## Supplementary Information for "The impact of whole genome duplications on the human gene regulatory networks"

### 1 Data Preprocessing

#### 1.1 Gene Nomenclature

The present study used data from many different sources, some of them produced in times pretty distant from one another. It was hence necessary to unify consistently the notation for the unique identifiers of the genes, in order to make them comparable in a sensible way. The Gene Symbol format was chosen, and all the genes appearing in the various datasets were translated to their official Gene Symbol, as indicated by the HGNC (HUGO Gene Nomenclature Committee - *https://www.genenames.org*). Gene Symbols can come in different possible statuses, and different actions were taken consequently:

- APPROVED: symbol left untouched;
- PREVIOUS: obsolete gene symbol, updated to the official one;
- WITHDRAWN: the gene symbol does not exist anymore, deleted;

For data available only in Ensembl ID format, we used the official mapping from one nomenclature to the other (obtained from the Ensembl BioMart interface - *https://www.ensembl.org/biomart*).

#### 1.2 Age distribution of paralogues

We show in fig. S1 that the dating of the pagarologues with their most recent common ancestor is consistent with the (independent) distinction between SSD and WGD couples and with the choice of considering SSD couples duplicated before *Sarcopterygii* as evolutionarily comparable to WGD couples. The figure also highlights the presence of SSD couples that were duplicated in relatively recent times, but which are a very small percentage of the total SSD couples. We did not consider such recent couples in all of the subsequent analyses.

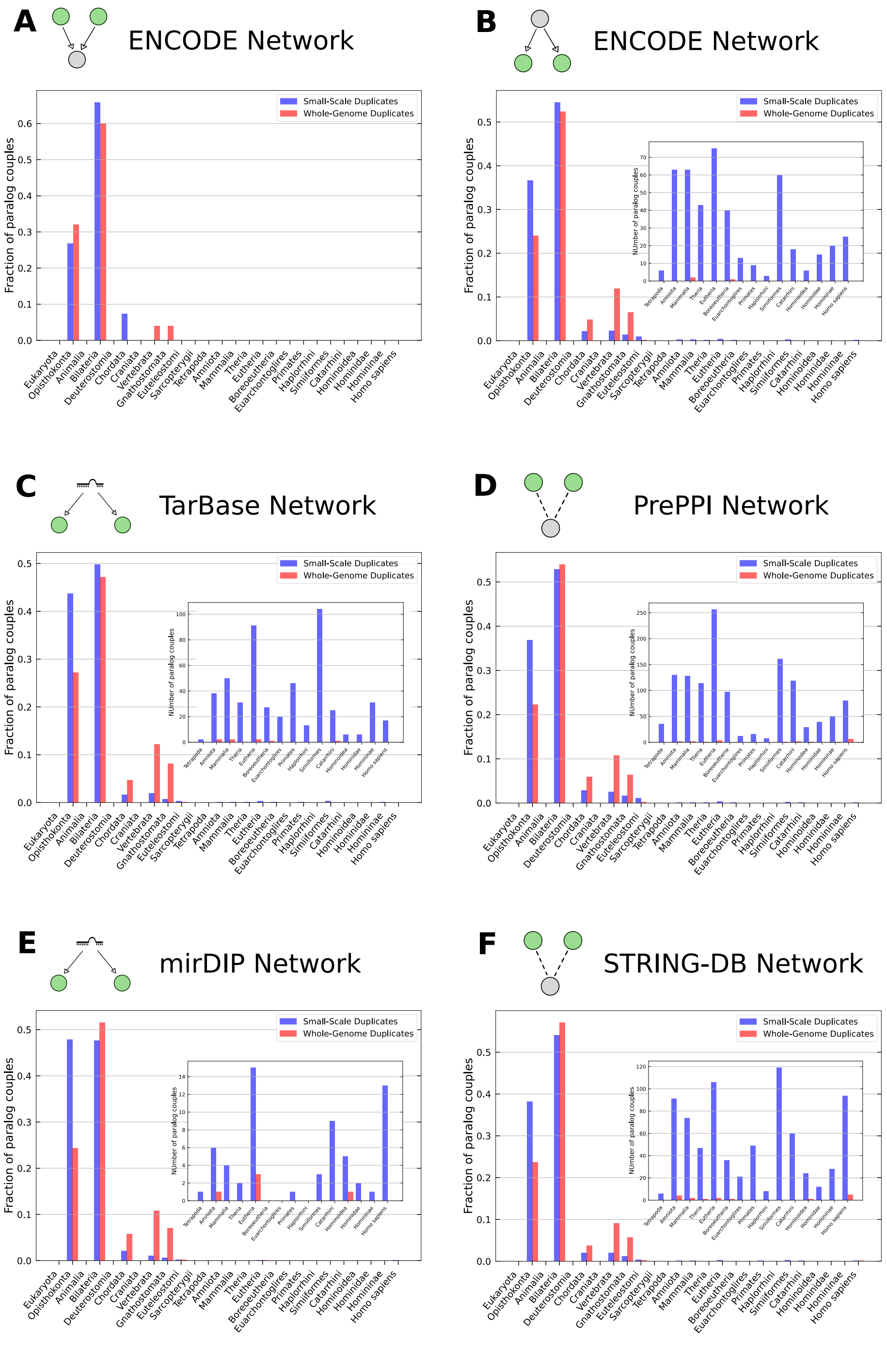

Figure S1: **Age distribution of WGD and SSD paralogues contained in the interaction networks.** The insets zoom in on the portion of the distribution after *Sarcopterygii*. The distribution for the ENCODE network is subdivided into TF couples **(A)** and target couples **(B)** for clarity.

#### 1.3 Effects of duplication age on interaction similarities

SSD couples are divided into *pre-Sarcopterygii* (comparable to WGD genes, used in all of the subsequent analyses) and *post-Sarcopterygii* (younger duplicates, discarded in the other analyses). We show in fig. S2 that younger duplicates indeed consistently show higher similarities with respect to older ones. Being their number very small, though, they do not introduce a significant bias in the analyses even when they are not discarded. Nonetheless, we eliminated them before proceeding with the analyses described in the paper, in order to keep the comparison with WGD genes as fair as possible.

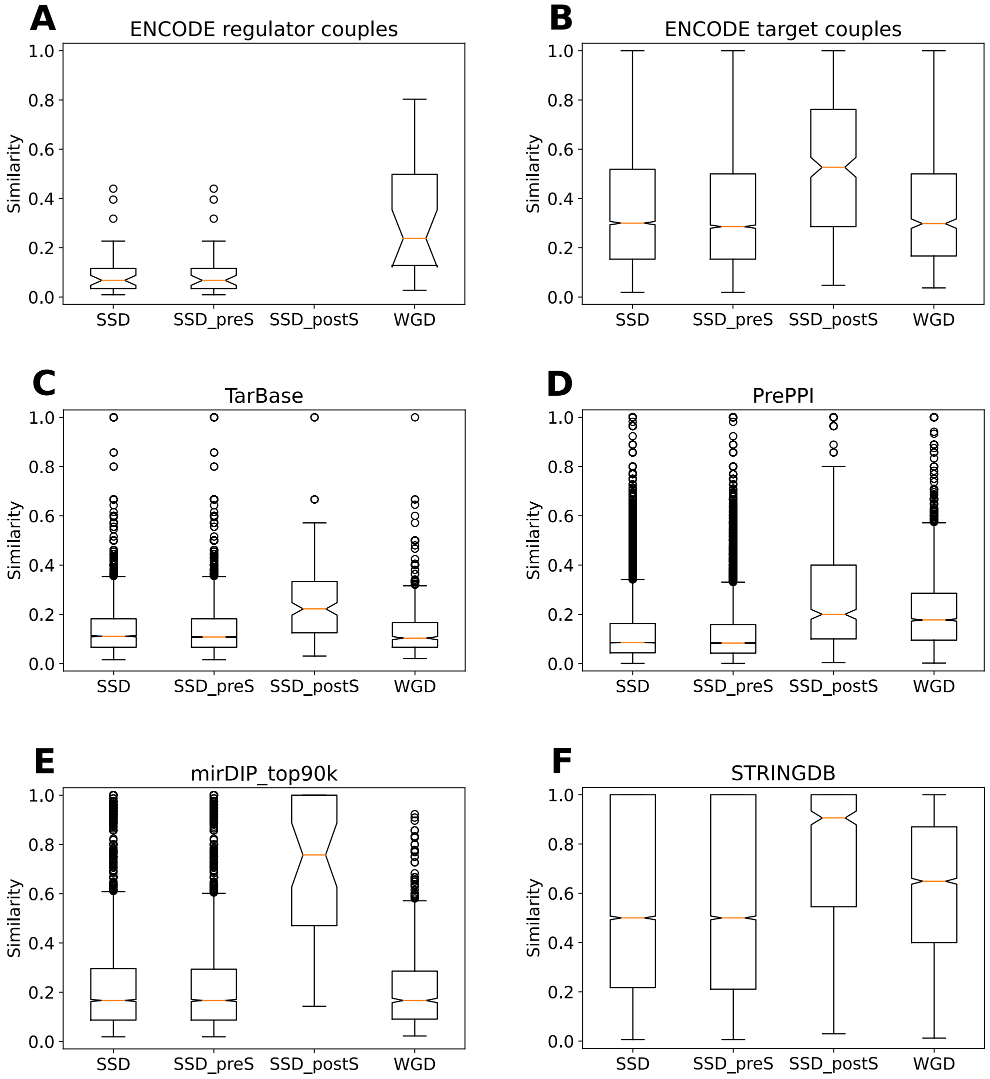

Figure S2: **Effects of different duplication ages on the similarity distributions.** The overall SSD distribution is marked as *SSD*, while *SSD_preS* represent older duplicates and *SSD_postS* represent younger duplicates. WGD couples are also shown for comparison.

### 2 Robustness of the results

#### 2.1 Degree distributions of the STRING and mirDIP networks

We show in this section that the degree distributions and the average degree of genes duplicated by SSD and WGD do not display any striking difference with respect to the global degree distributions, also in the case of the mirDIP and STRING datasets. Therefore, duplications do not display specific biases in terms of gene degree in the different networks considered. As explained in the main text, this is an important preliminary observation since the network motifs' enrichments might depend on the degree of the nodes.

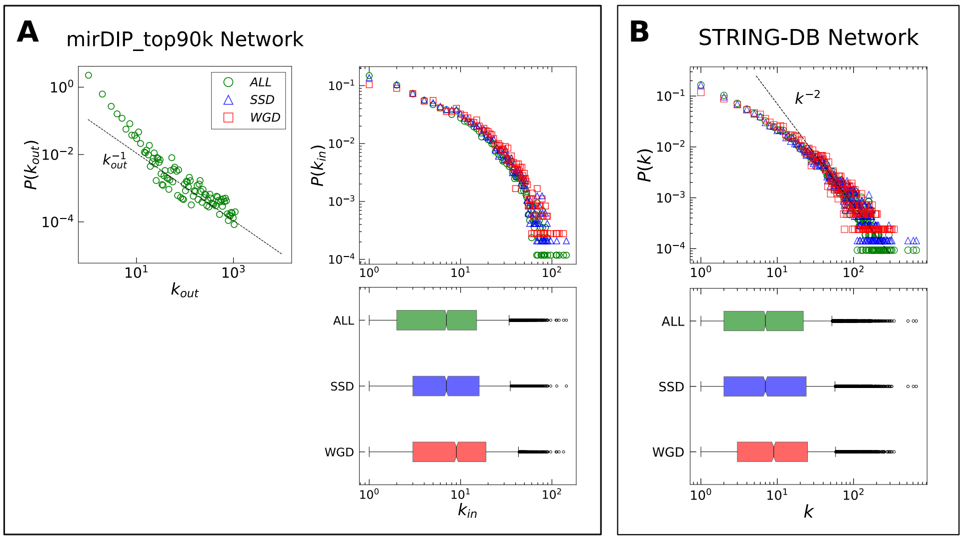

Figure S3: **Degree distributions**. Indegree ($k_{in}$) and outdegree ($k_{out}$) distributions of **(A)** the mirDIP miRNA-gene regulatory network and the degree (k) distribution of **(B)** the STRING protein-protein interactions network. Each degree distribution is shown both as a probability distribution (upper figure) and as a boxplot (lower figure). The global degree distribution of each network is represented in green, while the degree distributions of genes involved in a SSD couple and in a WGD couple are represented in blue and red, respectively. Dotted lines, corresponding to the reported scaling of the degree, are not the result of a fit and are shown as a reference only.

#### 2.2 Motif enrichment in the STRING network

We also show below that the results presented in the paper with the PrePPI network are independently confirmed by executing the same analyses on the STRING PPI network. In fact, the results for the paralogue couples which also have a protein-protein interaction are almost identical in the PrePPI and in the STRING case (see fig. 3 in the paper and fig. S4 below). The effects of paralogy relations in the similarity distribution of common PPI contacts, instead, are even more pronounced in the STRING data (fig. 4 in the paper and fig. S5 below).

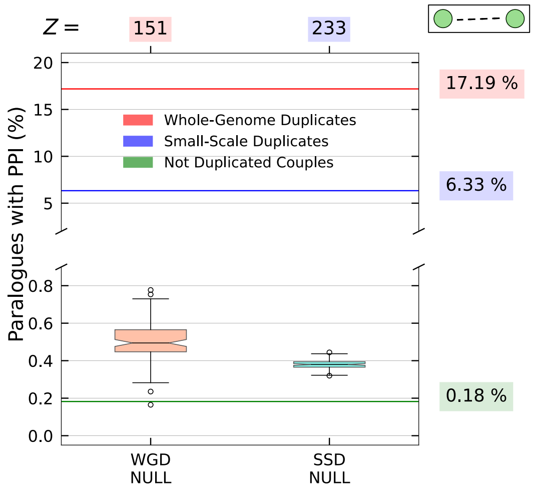

Figure S4: **Interactions of duplicated genes at the protein level (STRING database).** The percentages of gene pairs that present an interaction in the STRING database are indicated by the bold horizontal lines and explicitly stated in the labels on the right. The null model distributions are reported in the boxplots and the corresponding Z-scores are shown at the top.

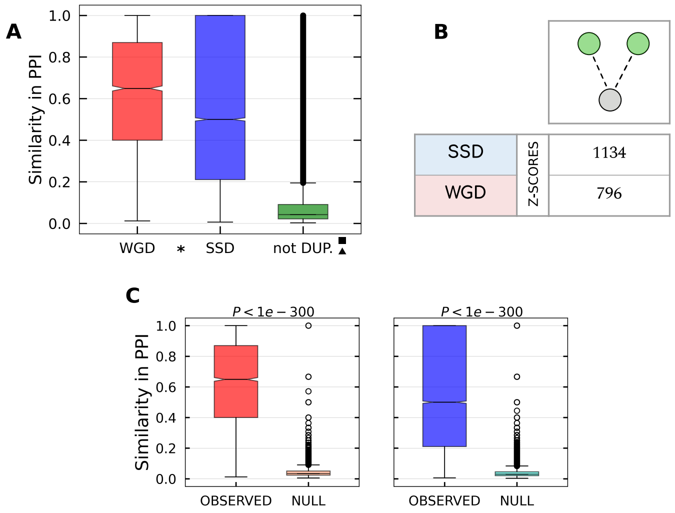

Figure S5: **Pairs of duplicated genes interacting with a third proteins (STRING database). (A)** Similarity distributions for WGD, SSD and not duplicated gene couples in the PrePPI network. All of the pairwise comparisons between distributions are statistically significant, as indicated by the presence of the symbols explained in the *Methods* section. **(B)** Z-scores measuring the enrichment of the co-interaction motif with respect to the null model. **(C)** Pairwise comparison between each real similarity distribution and the null distribution for the respective duplication type.

#### 2.3 Motif enrichment in the mirDIP network

Reported in fig. S6 are the results obtained for the similarity in regulators (that is, for the $\Lambda$ motif configuration) with the mirDIP miRNA-gene interaction network. The results shown are fully compatible with those presented in the paper in fig. 6.

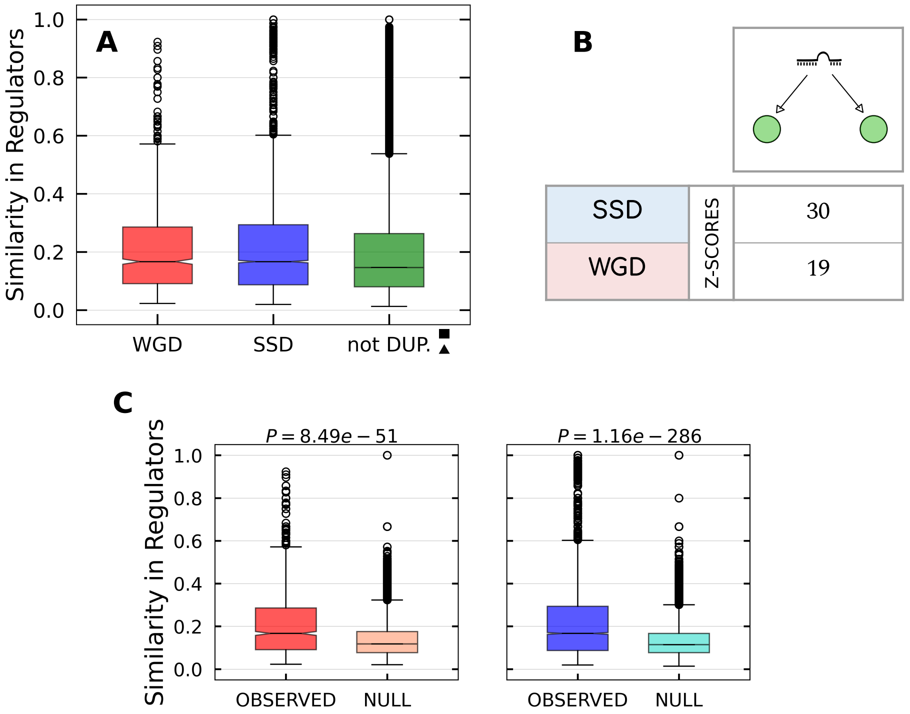

Figure S6: **miRNA $\Lambda$ motifs (mirDIP database)**. **(A)** Similarity distributions for WGD, SSD and not duplicated target genes couples in the mirDIP network. As indicated by the presence of the symbols explained in the *Methods* section, the difference between SSD and WGD distributions is not statistically significant, while both of them are significantly greater that the similarity distribution of non duplicated genes. **(B)** Z-scores measuring the enrichment of the $\Lambda$ motif with respect to the null model. **(C)** Pairwise comparison between each real similarity distribution and the null distribution for the respective duplication type.

#### 2.4 Alternative null models for mixed-type motifs

In fig. S7 we show the motif enrichments of mixed-type network motifs, for different choices of null models and databases. We observe that, despite some differences in the numbers, the overall trend is well consistent with the results reported in the main text.

**A** **ENCODE - PrePPI - TarBase**

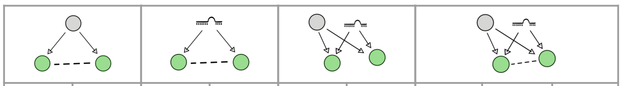

| | | ENCODE | PrePPI | TarBase | PrePPI | ENCODE | TarBase | ENCODE | TarBase | PrePPI |
|---|---|---|---|---|---|---|---|---|---|---|
| SSD | Z-SCORES | 21 | 47 | 4.6 | 46 | 17 | 12 | 3.6 | 0.6 | 18 |
| WGD | Z-SCORES | 12 | 31 | 4.6 | 47 | 1.0 | 0.2 | 6.9 | 2.5 | 5.3 |

**B** **ENCODE - STRING - TarBase**

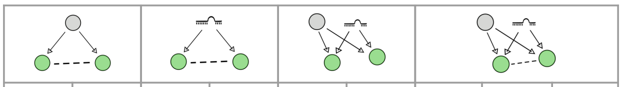

| | | ENCODE | STRING | TarBase | STRING | ENCODE | TarBase | ENCODE | TarBase | STRING |
|---|---|---|---|---|---|---|---|---|---|---|
| SSD | Z-SCORES | 21 | 37 | 3.8 | 68 | 17 | 12 | 7.6 | 6.4 | 14 |
| WGD | Z-SCORES | 9.8 | 31 | 3.5 | 39 | 1.0 | 0.2 | 6.0 | 2.2 | 6.7 |

**C** **ENCODE - PrePPI - mirDIP**

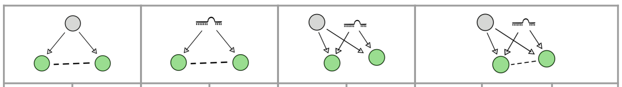

| | | ENCODE | PrePPI | mirDIP | PrePPI | ENCODE | mirDIP | ENCODE | mirDIP | PrePPI |
|---|---|---|---|---|---|---|---|---|---|---|
| SSD | Z-SCORES | 21 | 47 | 1.3 | 41 | 11 | 5.5 | 4.2 | 2.3 | 13 |
| WGD | Z-SCORES | 12 | 31 | 7.0 | 71 | 5.2 | 2.3 | 5.3 | 1.5 | 12 |

**D** **ENCODE - STRING - mirDIP**

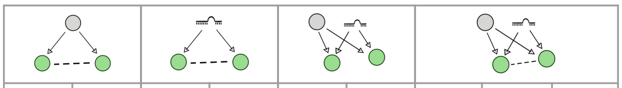

| | | ENCODE | STRING | mirDIP | STRING | ENCODE | mirDIP | ENCODE | mirDIP | STRING |
|---|---|---|---|---|---|---|---|---|---|---|
| SSD | Z-SCORES | 21 | 37 | 0.6 | 55 | 11 | 5.5 | 3.9 | -1.5 | 11 |
| WGD | Z-SCORES | 9.8 | 31 | 7.0 | 60 | 5.2 | 2.3 | 5.2 | 0.8 | 20 |

Figure S7: **Motifs with mixed-type regulatory interactions (different null models and network combinations).** Each table is referred to a different combination of databases, always comprising transcriptional, miRNA-gene and protein-protein interactions. In each table, from left to right: Transcriptional Δ motifs, miRNA-mediated Δ motifs, Mixed Bifan motifs, in which A pair of target genes are regulated both by a common TF and a common miRNA, and Mixed Bifan motif in which the two target genes interact at the protein level. The Z-scores are referred to the randomization of the network indicated in the column header. The table shown in the main text is a subset of **(A)**.
